# The BMP2/4 ortholog Dpp functions as an inter-organ signal that regulates developmental timing in *Drosophila*

**DOI:** 10.1101/180562

**Authors:** Linda Setiawan, Alexis L. Woods, Iswar K. Hariharan

## Abstract

For many organisms, developmental transitions are triggered by a neuroendocrine axis and are contingent upon multiple organs achieving sufficient growth and maturation. How the status of peripheral organs is communicated to the neuroendocrine axis is not known. In *Drosophila* larvae, metamorphosis is triggered by the steroid hormone ecdysone, secreted by the prothoracic gland (PG). Here we show that the BMP2/4 ortholog Dpp, which regulates growth and patterning of larval imaginal discs, also functions as a systemic signal to regulate developmental timing. Dpp from peripheral tissues, mostly imaginal discs, can reach the PG and inhibit ecdysone biosynthesis. As the discs grow, Dpp signaling decreases in the PG, thus alleviating the inhibition of ecdysone biosynthesis, and permitting entry into metamorphosis. We suggest that if a tissue can trap more morphogen locally as it grows and matures, then circulating levels of morphogen can provide a systemic readout of organ size and maturation.

**One Sentence Summary:** Dpp functions as a long-range endocrine signal between peripheral tissues and the prothoracic gland to regulate developmental timing in *Drosophila*.

---

Organismal development is often orchestrated by a neuroendocrine axis, such as the hypothalamic-pituitary axis in mammals. Mechanisms likely exist by which the growth and maturation of peripheral organs are monitored by the neuroendocrine axis prior to important developmental transitions. These mechanisms remain undefined. The onset of metamorphosis in *Drosophila*, a dramatic developmental transition, results from a steep increase in the level of the steroid hormone ecdysone (*1*) made by the prothoracic gland (PG), a part of the ring gland (Fig. 1A). The onset of pupariation in larvae is contingent upon sufficient growth (*2*), notably of the imaginal discs (*3*), the precursors of adult structures such as the wing. Regulators of ecdysone production in the PG include the peptide prothoracicotropic hormone (PTTH) (*4*), insulin signaling (*5, 6*), the activin pathway (*7*), the *bantam* (*ban*) microRNA (*8*), and under conditions of starvation, circulating Hedgehog made by enterocytes (*9*). It is not understood how growth of peripheral tissues can influence these pathways. The insulin/relaxin family member Dilp8, secreted by damaged or overgrown imaginal discs, delays pupariation (*10, 11*), but its role in regulating developmental timing under normal conditions is unclear.

**Figure 1.**
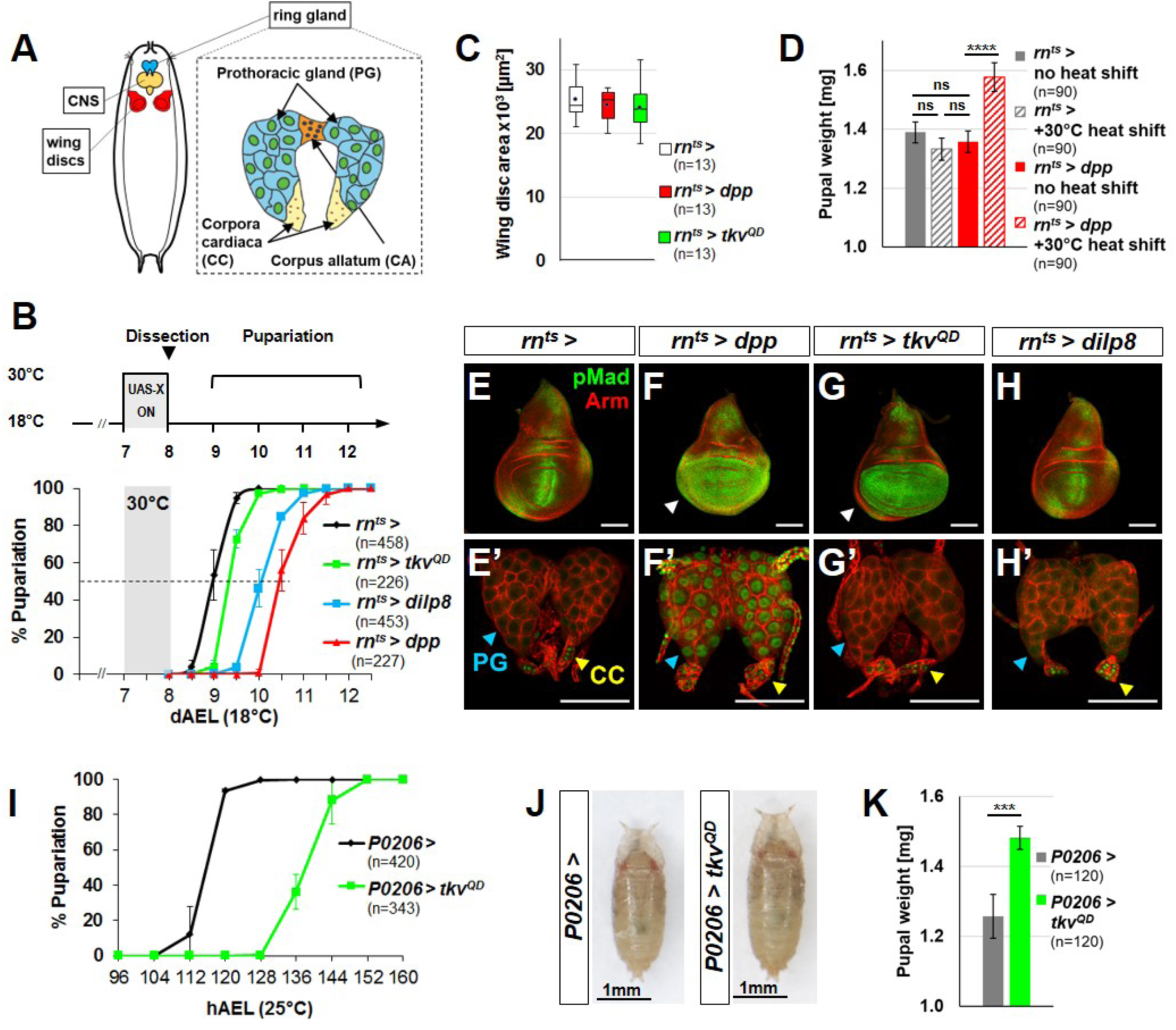
Expression of Dpp in peripheral tissues induces Dpp signaling in the PG and delays pupariation. (**A**) Schematic of ring gland (**B**) *rn*^*ts*^*>dpp, rn*^*ts*^*>tkv*^*QD*^, and *rn*^*ts*^*>dilp8* have a pupariation delay of 36.5 ± 3.5 h, 9.5 ± 3.0 h, and 26.0 ±3.7 h respectively (**C**) wing disc areas (**D**) pupal weights (**E-H**) wing discs (**E’-H’**) ring glands. Nuclear pMad is observed in the CC in all genotypes. (**I**) *P0206>tkv*^*QD*^ has a 22.2±2.1h developmental delay (**J**) pupal size (**K**) pupal weight increased in *P0206>tkv*^*QD*^ by 18%. Error bars indicate standard deviations, except (**C**) which is a box and whiskers plot. ns = not significant; *** p<0.001; **** p<0.0001. Scale bars = 100 μm.

The BMP2/4 ortholog, Dpp, functions as a morphogen to regulate growth and patterning within many tissues including imaginal discs (*12*). We increased *dpp* expression briefly during the early third larval instar (L3) using *rn-Gal4* and a temperature-sensitive repressor, Gal80^ts^ (hereafter *rn*^*ts*^>*dpp*) (Figure 1B)*. rn-Gal4* is expressed in wing discs and also some other tissues (Fig. S1A). *rn*^*ts*^*>dpp* did not increase wing disc size (Figure 1C) or adult wing size (not shown) but markedly delayed pupariation (Fig. 1B) resulting in larger pupae, likely due to an extended growth phase (Fig. 1D). Surprisingly, an activated form of the Dpp receptor Thickveins (Tkv^QD^) (*13*), (*rn*^*ts*^*>tkv*^*QD*^), which functions cell autonomously, elicited only a modest delay (Fig. 1B-C). Expression of *dpp*, but not *tkv*^*QD*^, increased Dpp signaling beyond the wing pouch (Fig. 1E-G, arrowheads in Fig. 1F, G) and in the PG, as assessed by increased nuclear pMad (Fig. 1E’-G’). *rn-Gal4* is not expressed in the ring gland but is expressed in the central nervous system, CNS (Fig. S1A). Delayed pupariation, nuclear localization of pMad in the PG, and increased pupal size all still occurred when neuronal expression of *rn-Gal4* was prevented (Fig. S1B-I). Thus, the Dpp that reaches the PG in these experiments is likely produced by more distant non-neuronal cells. Indeed, *dpp* expressed in various peripheral tissues delays pupariation (Fig. S2A-G) suggesting effects of Dpp outside its tissue of origin.

Did *dpp* delay pupariation indirectly via Dilp8 production? *rn*^*ts*^*>dpp* or *rn*^*ts*^*>tkv*^*QD*^ caused little cell death or *dilp8* expression (Fig. S3A-F) and *rn*^*ts*^*>dpp* delayed pupariation more than *rn*^*ts*^*>dilp8* (Fig. 1B) suggesting that *dpp* does not function upstream of *dilp8*. Conversely, *rn*^*ts*^*>dilp8* did not affect nuclear pMAD in the wing disc or PG (Fig. 1H, H’) indicating that Dilp8 does not activate Dpp signaling. Most importantly, *rn*^*ts*^*>dpp* delayed pupariation in a *dilp8* mutant implying that Dpp functions independently of *dilp8* (Fig. S3G). Finally, expressing *tkv*^*QD*^ using the ring gland driver *P0206-Gal4* (*5*) delayed pupariation and increased pupal mass (Fig. 1I-K). Pupariation was delayed using a PG-specific driver, but not using drivers expressed in either the corpus allatum (CA) or corpora cardiaca (CC) (Fig. S4A-B). Conversely, overexpression of *brinker* (*brk*) in the PG, which represses many Dpp target genes, arrested larvae in L2, yet caused wandering (prepupal) behavior, similar to the precocious metamorphosis observed with increased Activin signaling (*7*) (Fig. S4C-N). Thus, Dpp signaling in the PG itself regulates the timing of developmental transitions.

Can Dpp reach the PG from peripheral tissues? When *GFP-dpp* (*14*) was expressed using *rn-Gal4* (*rn*^*ts*^*>GFP-dpp*), GFP and nuclear pMad were detected in the PG but not in the immediately adjacent CA (Fig. 2A-B) suggesting localization dependent on binding to specific receptors. A different secreted protein ANF-GFP (*15*) was not detected in the PG (Fig. 2C-D). A processed form of GFP-Dpp was detected in the hemolymph of *rn*^*ts*^*>GFP-dpp* larvae (Fig. 2E). *Ex vivo*, Dpp can diffuse from discs to the PG. Nuclear pMad was not observed in the PG portion of ring glands cultured alone (Fig. 2F) but was observed in ring glands co-cultured with *rn*^*ts*^*>dpp* wing discs, or with *rn*^*ts*^*>dpp* larval hemolymph (Fig. 2G, H). Thus, Dpp can diffuse from discs to the PG via the hemolymph.

**Figure 2.**
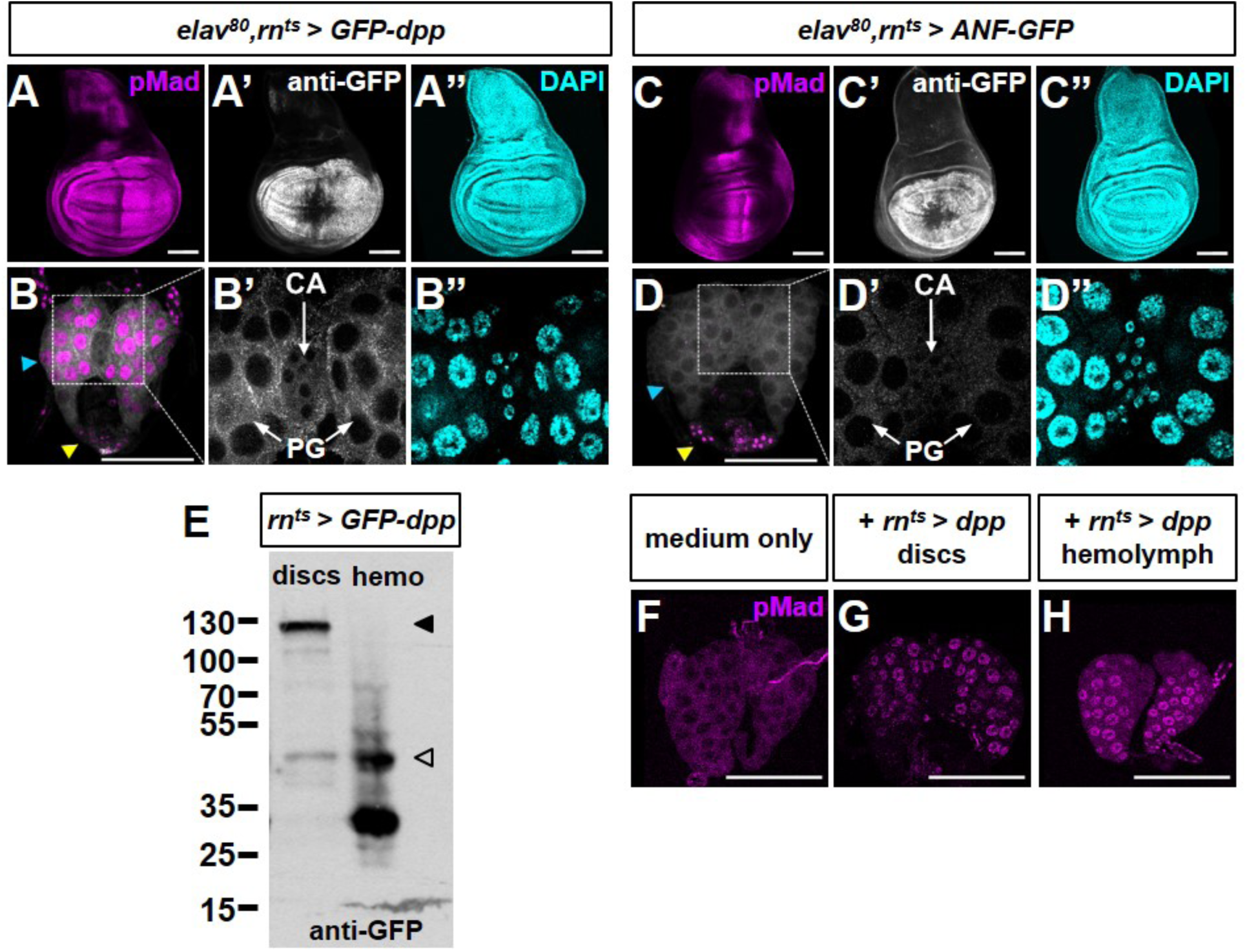
Dpp can diffuse from peripheral tissues to the PG via the hemolymph. *elav-Gal80 rn*^*ts*^*>GFP-dpp* wing disc (**A-A”**) and ring gland (**B-B”**). Note GFP in PG but not CA (**B’**). Blue arrowhead PG, yellow arrowhead CC. (**C-C”, D-D”**) *elav-Gal80 rn*^*ts*^*>ANF-GFP* wing disc (**C-C”**) and ring gland (**D-D”**). (**E**) Western blot of *rn*^*ts*^*>GFP-dpp* discs and hemolymph. Predicted unprocessed (filled arrowhead) and processed (unfilled arrowhead) forms. (**F-H**) Cultured ring glands.

Neither inducing apoptosis in the wing disc, nor disc overgrowth in *discs large* (*dlg*) hemizygotes resulted in nuclear pMad in the PG but caused local Dilp8 production (Fig. S5) and delayed pupariation (not shown). Thus, unlike Dilp8, Dpp is not released by discs in response to tissue damage or overgrowth.

To address a role for Dpp signaling in the PG during normal development, we extended our analysis to include ring glands at earlier time points during the final larval stage (L3) by visualizing nuclear pMad (Fig. 3A-C) and two reporters, *dad-RFP* (Fig. 3D-I) and *brk-GFP* (Fig. 3D-F, J-L). *dad-RFP* expression increases with Dpp signaling while *brk-GFP* expression increases as Dpp signaling decreases. In contrast to the PG in late L3 (120 h after egg lay, AEL), we observed nuclear pMad in the PG in early L3 (72 h AEL). As larvae progress through L3, Dpp signaling progressively decreases. By comparison, Dpp signaling was consistently high in the CC (Fig. 3A-C) and consistently low in the CA (Fig. 3J-L); its function in the CA and CC is not known.

**Figure 3.**
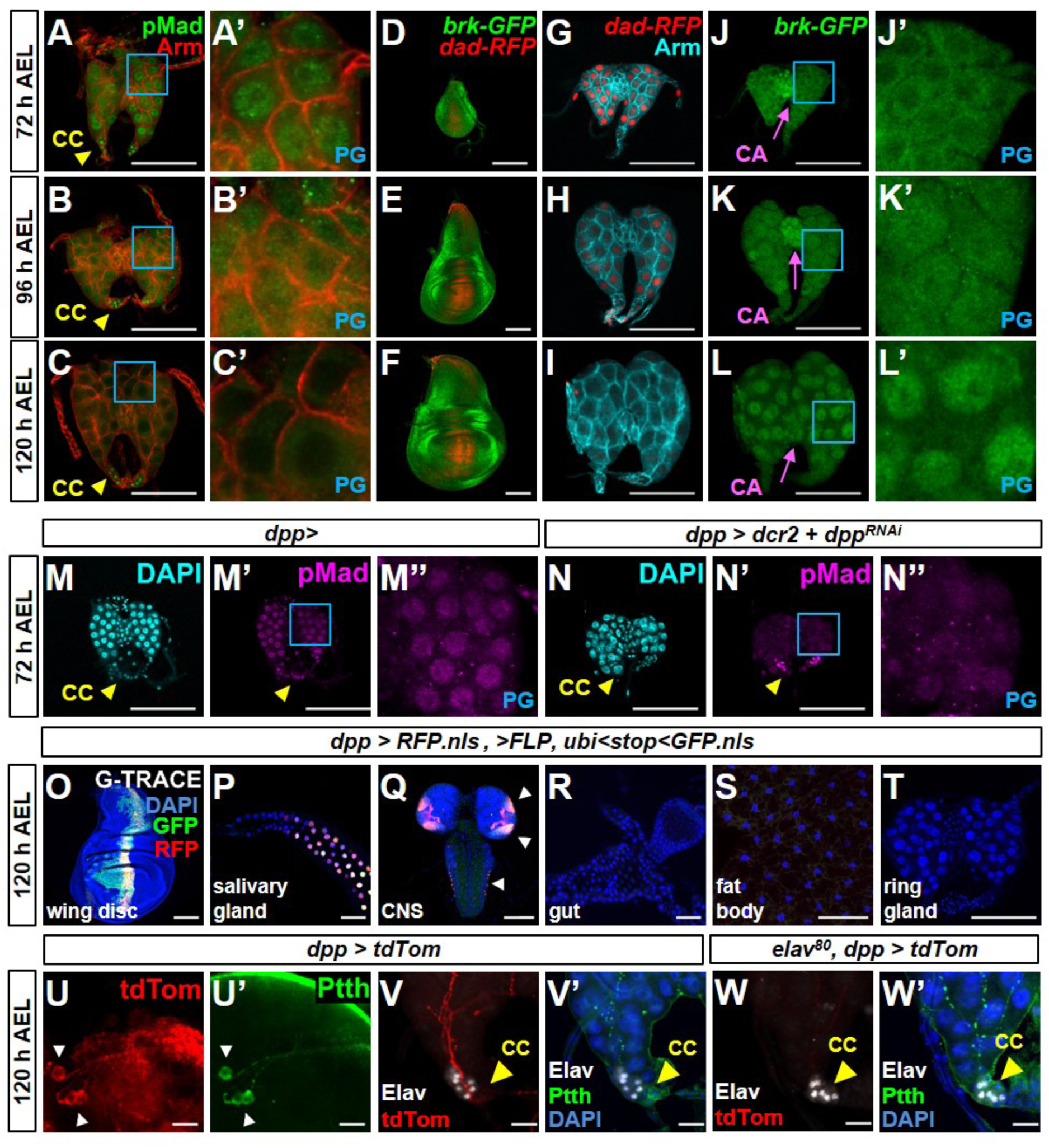
Dpp signaling in the PG decreases during L3. (**A-L’**) Ring glands and wing discs, 72, 96 and 120 h after egg lay (AEL) at 25°C. Boxed region in (**A-C**) at higher magnification in (**A’-C’**). (**D-F**) Wing discs expressing *dad-RFP* and *brk-GFP.* (**G-L’**) Ring glands with *dad-RFP* (**G-I**) or *brk-GFP* (**J-L**). Boxed regions at higher magnification in (**J’-L’**). (**M-N”)** Ring glands of indicated genotypes (**O-T**) *dpp-Gal4* expression analyzed using G-TRACE. Current (RFP) or prior (GFP) expression. (**U-V’**) Expression of *dpp-Gal4* in the PTTH-secreting neurons showing cell bodies (**U, U’**) and axons that innervate the PG (**V, V’**). (**W, W’**) *dpp-Gal4* expression in PTTH neurons blocked with *elav-Gal80*. Scale bars 100 μm except (**U-W’**) where they are 20 μm.

To help identify the source of Dpp that reaches the PG, we expressed *UAS-dpp*^*RNAi*^ using Gal4 drivers and examined pMad levels in the PG at 72 h AEL when nuclear pMad is normally observed. In *dpp-Gal4 UAS-dpp*^*RNAi*^ larvae, the level of nuclear pMad is greatly reduced (Fig. 3M-N). Thus, the Dpp must be produced by cells with current or past expression of *dpp-Gal4. dpp-Gal4* is expressed in some, but not all, Dpp-producing cells. G-TRACE (*16*) identifies both current (RFP) and previous (GFP) expression of *dpp-Gal4* (Fig. 3O-T); we detected expression in imaginal discs (Fig. 3O, Fig. S6A-C), the salivary glands (Fig. 3P) and CNS (Fig. 3Q) but not in the gut (Fig. 3R), fat body (Fig. 3S), ring gland (Fig. 3T) (also see legend to Fig. S6) or lymph gland (Fig. S6D). *dpp-Gal4* is expressed in the salivary glands (Fig. 3P) but *dpp* itself is not, either in embryos (*17*) or in wandering L3 larvae (*18*). In addition to the optic lobes and parts of the ventral nerve cord, CNS *dpp-Gal4* expression included the PTTH-expressing neurons which innervate the PG (Fig. 3U-V) and expression is blocked in the presence of *elav-Gal80* (Fig. 3W). However, expression of *GFP-dpp* in PTTH-producing neurons does not reach the PG nor is nuclear pMad observed in PG cells when *GFP-dpp* is expressed using *ptth-Gal4* (Fig. S6E-H). Additionally, expression of *dpp*^*RNAi*^ either using *elav-Gal4* or *ptth-Gal4* does not reduce pMad expression in the PG (Fig. S6I-V), indicating that the Dpp is unlikely to be from the PTTH neurons or other neurons. Taken together, these experiments suggest that the Dpp that reaches the PG is mostly from the imaginal discs. Additionally, the ability of *dpp-Gal4, UAS-dpp*^*RNAi*^ to reduce pMad levels in the PG indicates that the pathway is indeed mostly being activated by Dpp and not other ligands such as Gbb (*19*).

We examined the expression of several ecdysone biosynthesis enzymes (*20*) in FLP-out clones with altered Dpp signaling (Fig. S7A). In early L3, when Dpp signaling is high, reducing Dpp signaling by overexpressing *dad* (*21*) increased expression of Disembodied (Dib) (Fig. 4A) or Shadow (Sad) (Fig. S7B). In late L3, with lower Dpp signaling, augmenting Dpp signaling with activated Tkv reduced expression of Dib (Fig. 4B), Sad, Spookier (Spok) and Phantom (Phm) (Fig. S7C-E). Delayed pupariation and increased pupal weight caused by increased Dpp signaling in the PG can be suppressed partially by providing the ecdysteroid 20-OH ecdysone (20E) (Fig. 4C-D). In the PG, increased Dpp signaling increased *bantam (ban)* expression (Fig. 4E). Since *ban* expression delays pupariation and is negatively regulated by insulin signaling (*8*), Dpp and insulin signaling likely converge upstream of *ban* (Fig. S8A, B, D-N). Dpp signaling also functions antagonistically to the Activin pathway (Fig. S8C-E, O) as has been observed in the wing disc (*22*).

**Figure 4.**
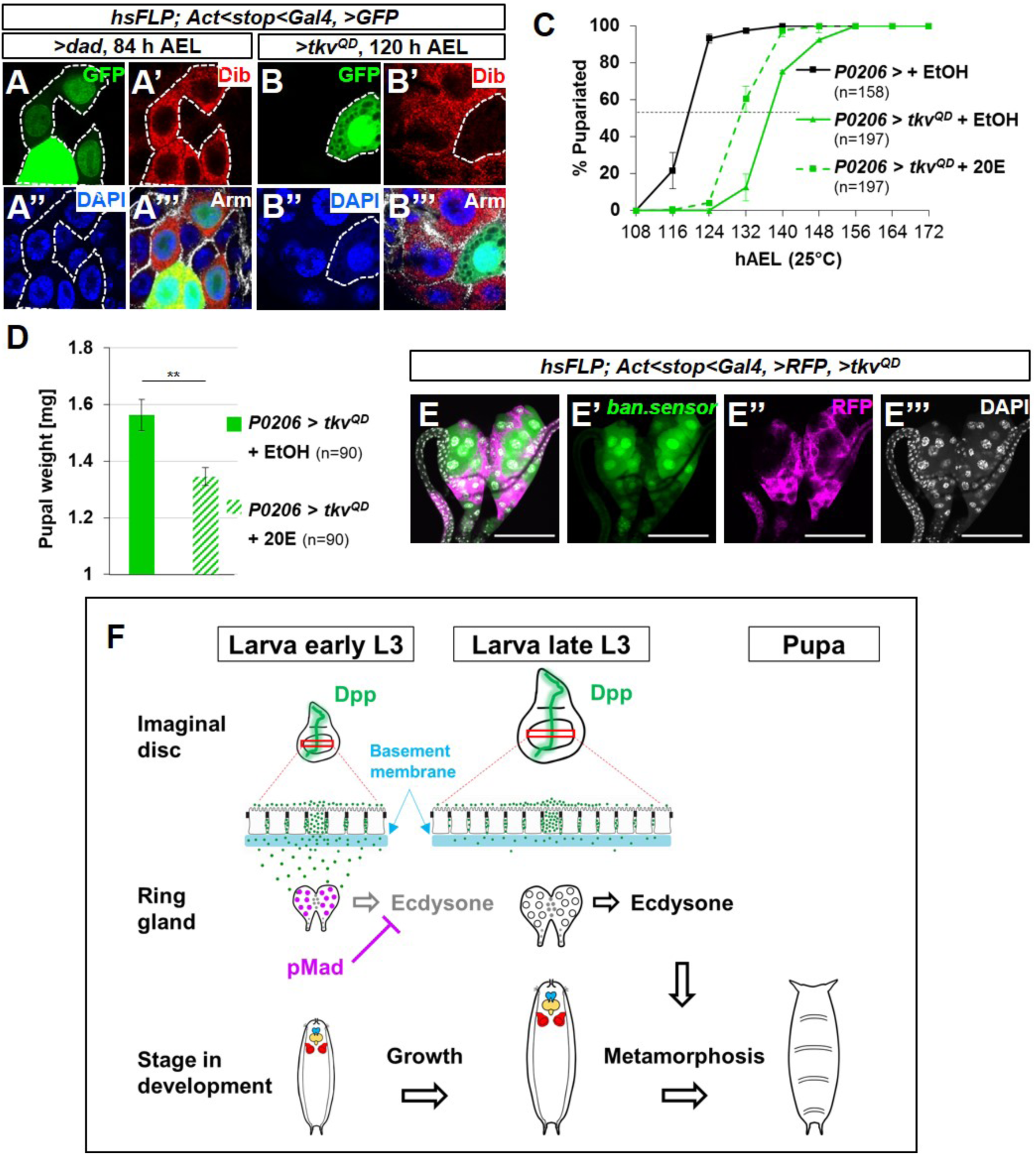
Dpp signaling in the PG represses ecdysone biosynthesis enzymes. (**A-B’’’**) FLP-out clones induced 48hAEL, ring glands dissected 84 h AEL or 120 h AEL, examined for Disembodied (Dib) expression. (**A-A’’’**) GFP-expressing cells express *UAS-dad*, lowering Dpp signaling. (**B-B’’’**) GFP-expressing cells express *UAS-tkv*^*QD*^, increasing Dpp signaling. (**C, D**) Effect of feeding 20-hydroxyecdysone (20E) to *P0206>tkv*^*QD*^ larvae. Mean pupariation times: *P0206>* +EtOH 119.0±0.5 h; *P0206>tkv*^*QD*^ + EtOH 136.8±0.3 h; *P0206>tkv*^*QD*^ 128.8±2.9 h (**C**) and pupal weight (**D**) ** p<0.01. (**E-E’’’**) Ring glands. FLP-out clones expressing *UAS-tkv*^*QD*^ and RFP. Reduced *ban* sensor expression (green in **E, E’**) indicates increased *ban* expression. Scale bars 100 μm. (**F**) Model for how discs influence Dpp signaling in the PG.

Dpp production, at least in wing discs, increases as discs grow during L3 (*23*). Dpp from discs can pass through the basement membrane into the hemolymph (*24, 25*). It is therefore surprising that Dpp signaling in the PG decreases concurrently. One possibility is that the PG becomes less sensitive to Dpp, perhaps by a downregulating Dpp receptors or other signaling components. However, pulses of Dpp in late L3 using *rn*^*ts*^*>dpp* elicit undiminished reporter responses in the PG (not shown). Although Dpp production in discs increases during L3 (*23*), disc growth results in a disproportionate increase in the number of cells that do not produce Dpp, yet can bind Dpp, thus expanding the “morphogen sink”. Additionally, extracellular matrix proteins continue to be deposited on the basement membrane throughout L3 (*26*). For either or both of these reasons, as discs grow, more Dpp might be retained within the discs and less allowed to diffuse into the hemolymph and reach the PG (model in Fig. 4F). Any increase in hemolymph volume would further dilute Dpp in the hemolymph. Thus, growth and maturation of the larva can influence the timing of pupariation. Dpp therefore has dual roles: one within tissues to regulate patterning and growth and one in inter-organ communication to regulate developmental timing. It will be of interest to determine if vertebrate BMP proteins also have dual functions.

## ACKNOWLEDGEMENTS

We thank many colleagues in the fly community for constructive suggestions, María Domínguez, Marcos González-Gaitán, Pierre Léopold, Michael O’Connor, Lynn Riddiford, Chris Rushlow, Michael Thomas Marr, Hilary Ashe and Konrad Basler for fly stocks and antibodies, the Bloomington, VDRC and TRiP stock centers, Octavio Bejarano for technical assistance and Jo Downes Bairzin, David Bilder, Robin Harris, Nipam Patel, Taryn Sumabat and Melanie Worley for comments on the manuscript. IKH was funded by NIH grants R01GM085576 and R35GM122490 and an American Cancer Society Research Professor Award (RP-16-238-06-COUN).

## SUPPLEMENTARY MATERIALS

## MATERIALS AND METHODS

### *Drosophila* strains and husbandry

Animals were raised on standard medium as used by the Bloomington Drosophila Stock Center. Fly stocks used are: *w*^1118^ as wild-type control for all experiments, *Oregon-R, rn-Gal4,tub-Gal80*^*ts*^*/TM6B-Gal80, rn-Gal4,tub-Gal80*^*ts*^*,UAS-egr/TM6B-Gal80* and *rn-Gal4,tub-Gal80*^*ts*^*,UAS-rpr/TM6B-Gal80* (*27*), *UAS-dpp/TM6B* (*19*), *UAS-tkv*^*Q*253*D*^*/TM6B* (*13*), *dad-nRFP/CyO* (*23*), *UAS-GFP-Dpp* (*14*), *UAS-dilp8::3xFLAG* (*11*), *dpp-LG* and *lexOEGFP:dpp/TM6B* (*28*), *UAS-bantamA* and *bantam.sensor* (*29*), *P0206-Gal4, phm-Gal4* and *ptth-Gal4* (gifts from Lynn Riddiford), *UAS-dad* and *UAS-brk* (gifts from Chris Rushlow), *UAS-babo*^*CA*^ (gift from Michael O’Connor), *elav-Gal80* (*30*) (gift from Lily Yeh Jan and Yuh-Nung Jan), *dlg*^40^-^2^ (gift from David Bilder). Stocks that were obtained from the Bloomington Drosophila Stock Center: *UAS-InR*^*ACT*^ (#8263),*dpp-Gal4/TM6B* (#1553) *UAS-dpp* (#1486), *UAS-tkv*^*CA*^ (#36537), *brk-GFP.FPTB* (#38629), *dilp8*^*MI*00727^ (#33079), *Aug21-Gal4* (#30137), *AKH-Gal4* (#25684), *UAS-dcr2* (#24650), *UAS-dpp RNAi* (#25782, #33618, #36779), *r4-Gal4* (#33832), *UAS-preproANF-EMD* (#7001), *G-TRACE-3* (#28281), *UAS-Mad*^*RNAi*^ (#40397), *UAS-Med*^*RNAi*^ (#19688), *AbdB-Gal4* (#55848), *UAS-tdTom* (#36328), UAS-tdGFP (#35836).

### Immunohistochemistry and microscopy

Larvae were dissected in PBS, fixed 20 min in 4% PFA, permeabilized with 0.1% Triton X-100, and blocked with 10% normal goat serum. Primary antibodies used are: rabbit anti-Smad3 (Abcam #52903, 1:500), rabbit anti-Sad (1:250), rabbit anti-Phm (1:250), rabbit anti-Dib (1:250) and guinea pig anti-Spok (1:1000) (gifts from Michael O’Connor), rabbit anti-Ptth (1:100) (gift from Pierre Léopold), mouse anti-Armadillo N2 7A1 (*31*)(DSHB, 1:100), mouse anti-Dlg 4F3 (*32*)(DHSB, 1:100), rabbit anti-GFP (Torrey Pines Biolabs #TP401, 1:500), mouse anti-GFP (Abcam AB290, 1:500), Cambridge, MA), rabbit anti-cleaved DCP-1 (Cell Signaling Asp216, 1:250), rabbit anti-FOXO (1:1000) (gift from Michael Thomas Marr) (*33*). Secondary antibodies used are: goat anti-mouse 555 (Invitrogen #A32727), goat anti-mouse 647 (Invitrogen #A32728), goat anti-rabbit 555 (Invitrogen #A32732), goat anti-rabbit 647 (Invitrogen #A32733) goat anti-guinea pig 555 (Invitrogen #A-21435) (all, 1:500) as well as phalloidin-TRITC (Sigma #P1951, 1:500) and DAPI (Invitrogen #D1306, 1:500). Samples were mounted in SlowFade Gold (Invitrogen #S36937) and imaged on a Zeiss 700 LSM confocal microscope.

### Developmental timing assay and *rn*^*ts*^*>* temperature shift experiments

Fertilized eggs were collected on grape juice plates for 4h. L1 stage larvae were transferred onto standard Bloomington food supplemented with yeast paste at a density of 50 animals per vial. For constitutive expression without the presence of a temperature-sensitive Gal80, animals were raised consistently at 25°C and pupal counts were taken every 8 h. Three independent experiments were conducted for each condition. *rn*^*ts*^*>* animals were raised at 18°C until day 7 (early third instar), then transferred to 30°C for a 24 h temperature shift and subsequently returned to 18°C. Pupal counts were taken every 12 h. Three independent experiments were conducted for each condition.

### 20-Hydroxyecdysone feeding

A stock solution of 10 mg/ml 20E (Sigma #H5142) in ethanol was prepared. 50 larvae per vial were raised on standard Bloomington food. At 72 h AEL, 1mg 20-Hydroxyexcdysone or (100μl stock solution or ethanol for controls) was added to each vial.

### Quantification of wing discs and pupal weight

Pupae from developmental timing assays were collected at the pharate adult stage were cleaned with 70% ethanol, dried and weighed in groups of 30 in three or four independent experiments. GraphPad Prism 6 was used to determine statistical significance between groups by one-way ANOVA using Tukey’s or Dunnett’s test. Pupae were placed on double-sided adhesive tape for imaging using a Leica transmitted light microscope (TL RCI, Germany). Adobe Photoshop was used to quantify the area of imaginal disc confocal images dissected from larvae from developmental timing assays.

### Hemolymph extraction

Larvae were bled into chilled PBS supplemented with protease inhibitor (Roche # 11697498001) on a cold aluminum block using a fine tungsten needle to puncture the cuticle.

### Western Blotting

For homogenates, staged larvae at 72 h AEL were transferred into chilled buffer containing 35mM Tris-HCl pH 6.8, 129mM NaCl, 4mM KCl, 2mM CaCl_2_ supplemented with protease inhibitor (Roche #11697498001). Larvae were homogenized for 1 min with a microtube homogenizer, followed by centrifugations for 5 min at 16,000g and 2 min at 16,000g at 4°C. Homogenates or hemolymph were boiled for 10 min in Laemmli sample buffer, run on 10% Mini-Protean TGX gels (Bio-Rad) and transferred to nitrocellulose membrane (Bio-Rad). Primary antibody used was rabbit anti-GFP (Torrey Pines Biolabs #TP401, 1:1000). Protein bands were detected with secondary antibody HRP anti-rabbit (Santa Cruz Biotechnology #sc-2030, 1:2500) and Western Lightning Plus-ECL (PerkinElmer #NEL103001EA).

### *Ex vivo* organ culture

20 brain-ring gland complexes were dissected from wandering third instar Oregon R larvae in Schneider’s medium (Gibco # 21720024) by pulling mouth hooks from which salivary glands, lymph gland and fat body was removed. Complexes were subsequently co-cultured with either wing imaginal discs or hemolymph from *rn*^*ts*^*>dpp* larvae larvae in Schneider’s medium supplemented with 10% FBS (Invitrogen #26140087) and Penicillin-Streptomycin at 1:100 of a 5,000 U/ml stock (Gibco #15070063).

### Irradiation

Density controlled third instar larvae were placed on shallow food plates and irradiated with 45 Gy in an X-ray cabinet (Faxitron, Tucson, AZ), followed by dissection after 12h.

### Heat-shock clone induction

Flies with *UAS* transgenes were crossed to either *ywhsFlp;;Act>>Gal4,UAS-GFP* or *ywhsFlp;;Act>>Gal4,UAS-RFP* and raised at 25°C. Larvae were staged and density controlled as described for developmental timing assay, then heat shocked in a 37°C water bath for 5 min at 24 or 48 hAEL before returning to 25°C until dissection.

**Supplementary Figure S1:**
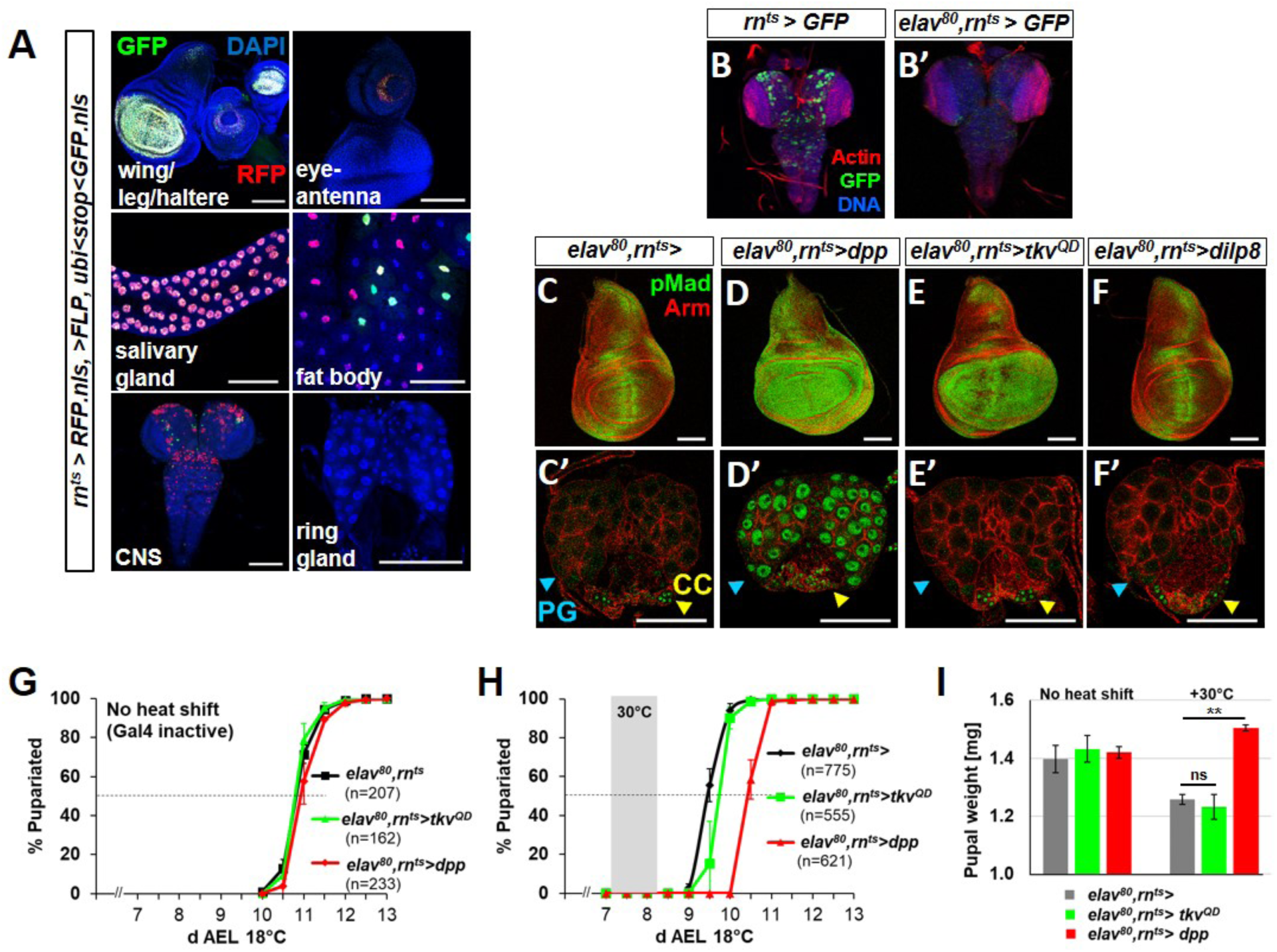
Characterization of *rn*^*ts*^*>* expression and investigation of the role of expression in the central nervous system (CNS) on pupariation delay. **(A)** Characterization of expression of *rn*^*ts*^*>* using G-TRACE (*16*). RFP (red) indicates recent expression, GFP (green) indicates either recent or past expression. Expression is observed in imaginal discs and the salivary glands, in scattered nuclei in the fat body and in the central nervous system (CNS) i.e. brain and ventral nerve cord. No expression was detected in any portion of the ring gland. (**B, B’**) Expression of *rn*^*ts*^*>GFP* in the CNS in the absence (**B**) or presence (**B’**) of *elav-Gal80* (*elav*^80^). (**C-F, C’-F’**) Effect of reducing CNS expression of *rn*^*ts*^*>* using *elav-Gal80* on Dpp signaling in wing discs (**C-F**) and the ring gland (**C’-F’**). Unlike in the PG (blue arrowhead), pMad is observed in the CC (yellow arrowhead) in all genotypes. (**G, H**) Timing of pupariation of different genotypes cultured at 18°C either without (**G**) or with **(H)** a shift to 30°C for 24 h beginning on day 7 AEL. Delays when compared to *elav*^80^*, rn*^*ts*^*>* are: *elav*^80^*, rn*^*ts*^*> dpp* 28.4±6.4 h; *elav*^80^*, rn*^*ts*^*> tkv*^*QD*^ 8.0±1.2 h **(I)** Pupal weights of the genotypes shown in (**G**) and (**H**). These are, without a heat shift, *elav*^80^ *rn*^*ts*^*>* 1.40±0.05 mg; *elav*^80^*, rn*^*ts*^*> tkv*^*QD*^ 1.43±0.05 mg; *elav*^80^*, rn*^*ts*^*> dpp* 1.42±0.02 mg (n=90 per genotype) and with a heat shift *elav*^80^ *rn*^*ts*^*>* 1.26±0.02 mg; *elav*^80^*, rn*^*ts*^*> tkv*^*QD*^ 1.23±0.04 mg; *elav*^80^*, rn*^*ts*^*> dpp* 1.50±0.01 mg (n=90 per genotype). Error bars indicate standard deviations. ns indicates not significant; ** indicates p < 0.01. All scale bars = 100 μm.

**Supplementary Figure S2:**
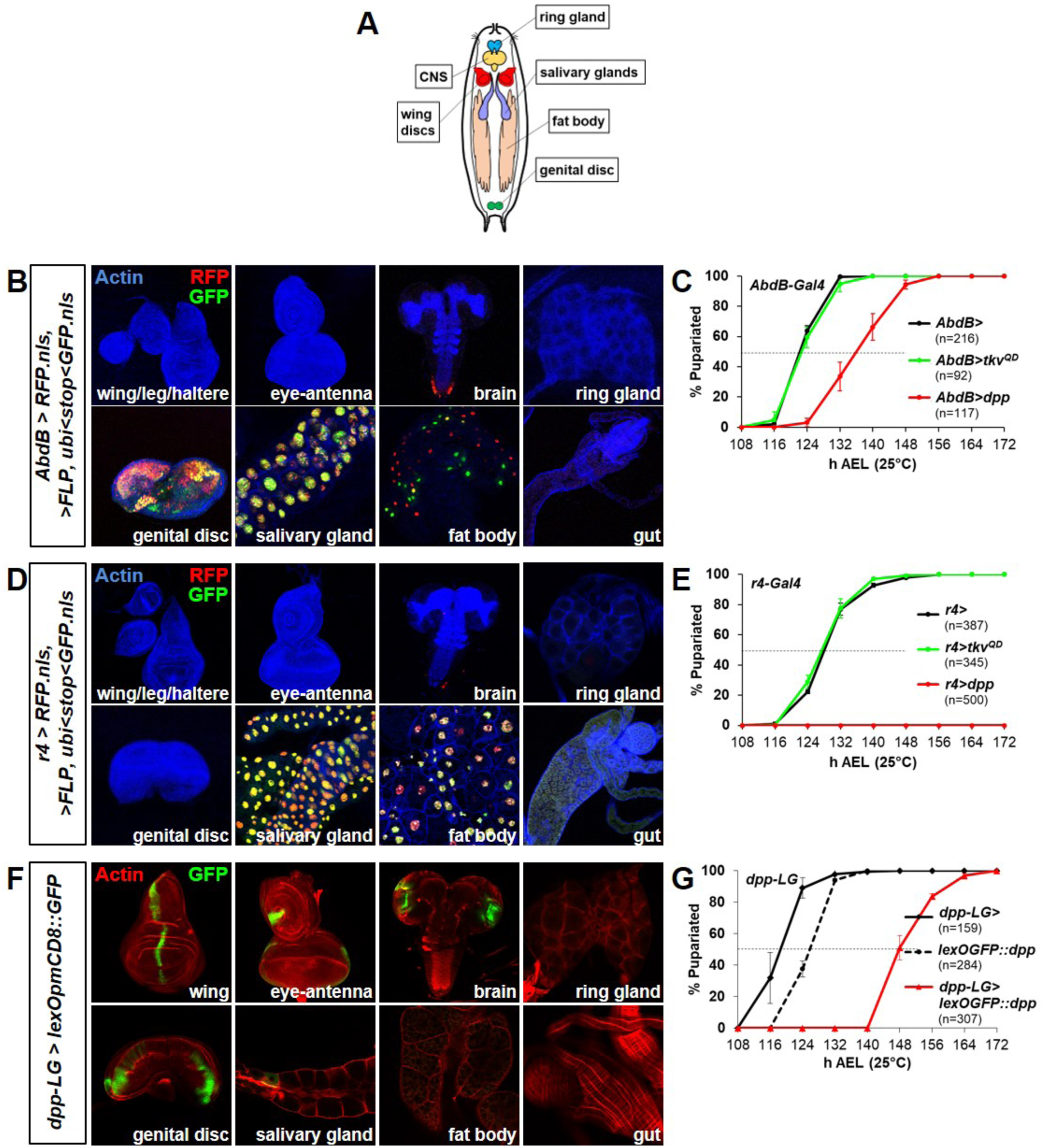
Effects of expression of *dpp* and *tkv*^*QD*^ in a variety of tissues on the timing of pupariation. (**A**) Cartoon of larva showing tissues relevant to experiments in (**B-G**) (**B-G**) Effect of expressing *dpp* or *tkv*^*QD*^ using heterologous expression systems on pupariation timing. The expression patterns of *AbdB-Gal4* (**B**), *r4-Gal4* (**D**) are shown using G-TRACE and of *dpp-LG* using a lexOp-mCD8::GFP (**F**). The corresponding graphs (**C, E, G**) show the effect on the timing of pupariation.

**Supplementary Figure S3:**
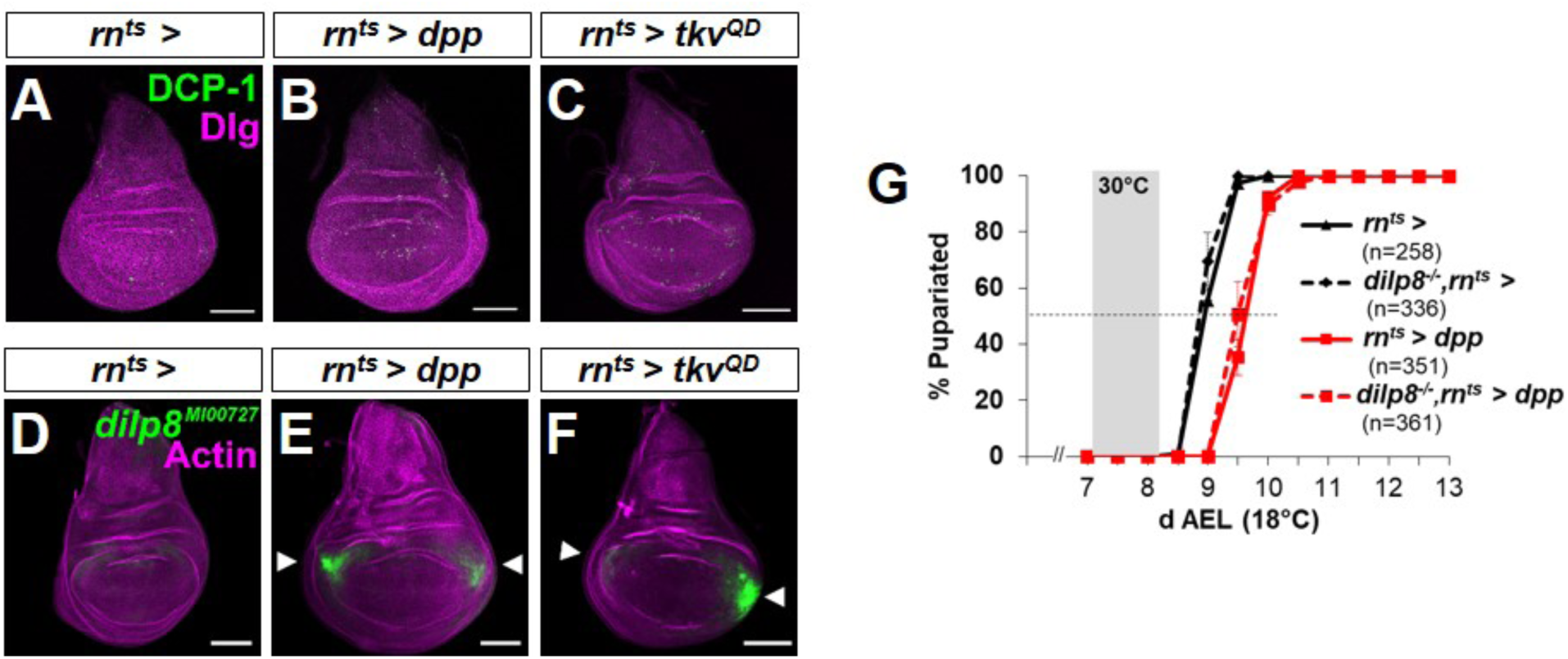
*rn*^*ts*^*>* expression of *dpp* or *tkv*^*QD*^ in wing discs causes relatively little cell death or *dilp8* expression. (**A-C**) Effect of increased expression of *dpp* or *tkv*^*QD*^ on levels of apoptosis (visualized by anti-DCP-1). Anti-Dlg labels cell outlines. (**D-F**) Effect of increased expression of *dpp* or *tkv*^*QD*^ on expression of a GFP gene trap in the *dilp8* gene (*11*). Arrowheads indicate increased *dilp8* expression in lateral regions of the wing pouch suggestive of mild overgrowth in this region. Phalloidin (actin) labels cells in the disc. Scale bars are 100 μm. (**G**) Comparison of the delay in pupariation elicited by *rn*^*ts*^*>dpp* in a wild-type and *dilp8* homozygous mutant background. The time course of pupariations of *rn*^*ts*^*>* (no *UAS-dpp*) in the two backgrounds are also shown. Mean pupariation times are *rn*^*ts*^*>* 8.93±0.03 d *; rn*^*ts*^*> dpp* 9.62±0.03 d AEL (delay = 16.4±0.3 h); *dilp8 -/-, rn*^*ts*^*>* 8.87±0.03 d; *dilp8 -/-, rn*^*ts*^*>dpp* 9.62±0.11d AEL (delay = 18.00±0.52 h).

**Supplementary Figure S4:**
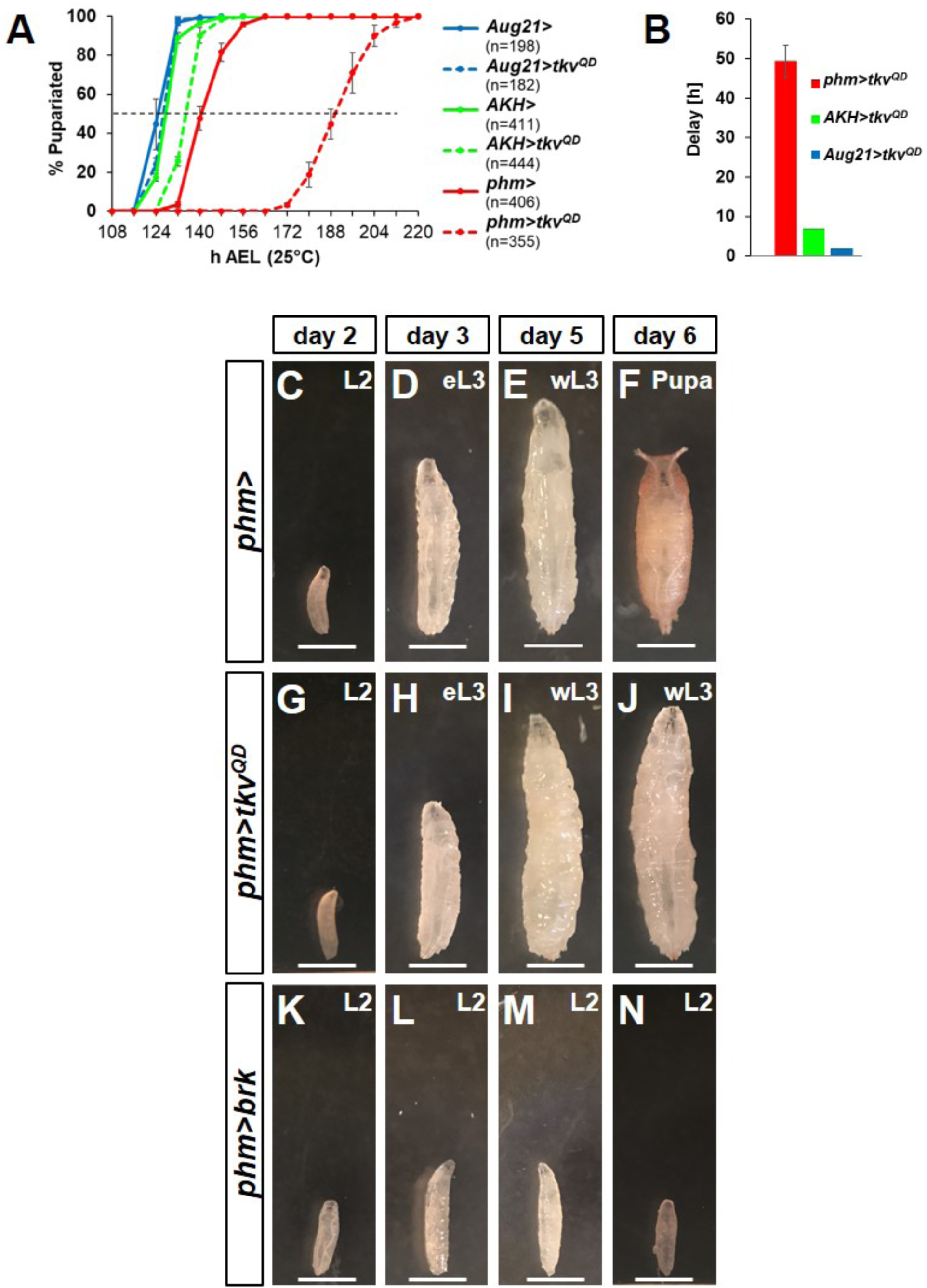
Effects of altering Dpp signaling or of *brinker* (*brk*) expression in the PG on pupariation. (**A, B**) Effect of expressing *tkv*^*QD*^ in different parts of the ring gland on pupariation timing. *phm-Gal4* drives expression in the PG (*34*); *AKH-Gal4* (*35*) is expressed in the CC; *Aug21-Gal4* is expressed in the CA (*36*). Mean pupariation times are: *phm*> 140.3±1.5 h; *phm>tkv*^*QD*^ 189.7±2.5 h; *Aug21*> 124.3±1.5 h; *Aug21>tkv*^*QD*^ 126.3±0.6 h; *AKH*> 127.8±0.3 h; *AKH>tkv*^*QD*^ 134.7±0.6 h The delays shown in (**B**) show the effect of including *UAS-tkv*^*QD*^ when compared to the corresponding Gal4 driver line alone. Error bars are standard deviations. (**C-N**) Temporal comparison of *phm>* (**C-F**)*, phm>tkv*^*QD*^ (**G-J**) and *phm>brk* (**K-N**) larvae. Images of stage- and density-controlled animals raised at 25°C. On day 2 AEL, (**C**) control *phm>* animals, (**G**) *phm>tkv*^*QD*^ and (**K**) *phm>brk* animals are at the second instar stage (L2). *phm*> and *phm>tkv*^*QD*^ animals progress to the early third instar (eL3) on day 3 (**D, H**) and wandering third instar (wL3) on day 5 (**E, I**). *phm>* animals pupariate by day 6 (**F**) while *phm>tkv*^*QD*^ animals remain in wL3 (**J**) until a much delayed pupariation on day 8 or day 9. *phm>brk* animals do not progress to eL3 on day 3 (**L**) and remain small and in second instar through days 5 and 6 (**M, N**). However, while still in L2, they exhibit wandering behavior for multiple days followed by lethality without pupariation. Wandering behavior in L2 is reminiscent of the early pupariation phenotype observed with expression of a constitutively-active form of the Activin receptor, Baboon (*7*). Scale bars are 1 mm.

**Supplementary Figure S5:**
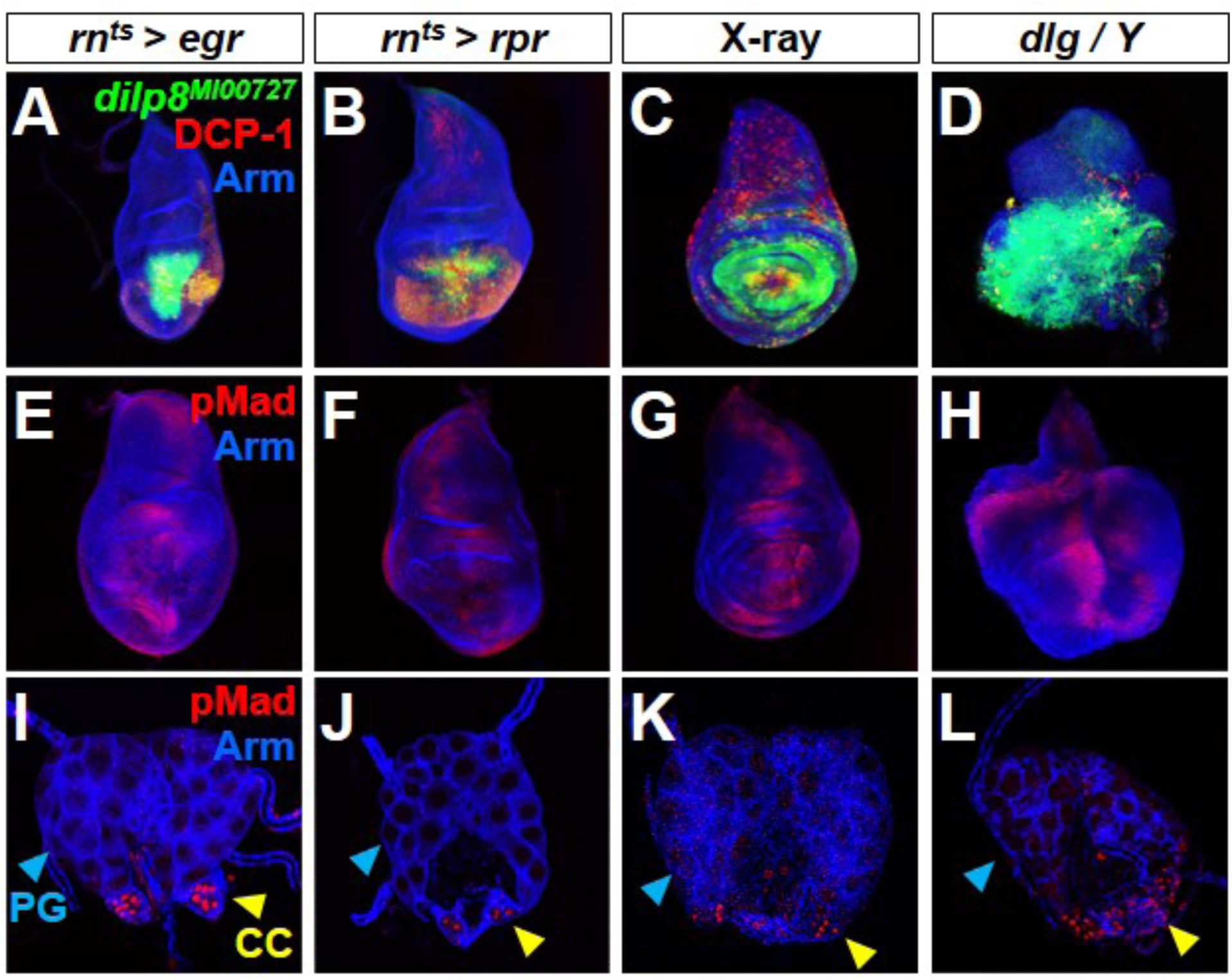
Cell death and disc overgrowth result in *dilp8* expression in discs but not in ectopic Dpp signaling in either the disc or ring gland. Wing discs (**A-H**) and ring glands (**I-L**) shown under conditions that increase cell death by expression of *eiger* (*egr*)(**A, E, I**), *reaper* (*rpr*) (**B, F, J**), or X-ray irradiation (**C, G, K**) or neoplastic overgrowth of the disc in *dlg* hemizygotes (**D, H, L**). Each of these conditions caused *dilp8* expression in the disc but did not result in ectopic nuclear pMad either in the wing disc or PG. The CC cells have nuclear pMad in all genotypes.

**Supplementary Figure S6:**
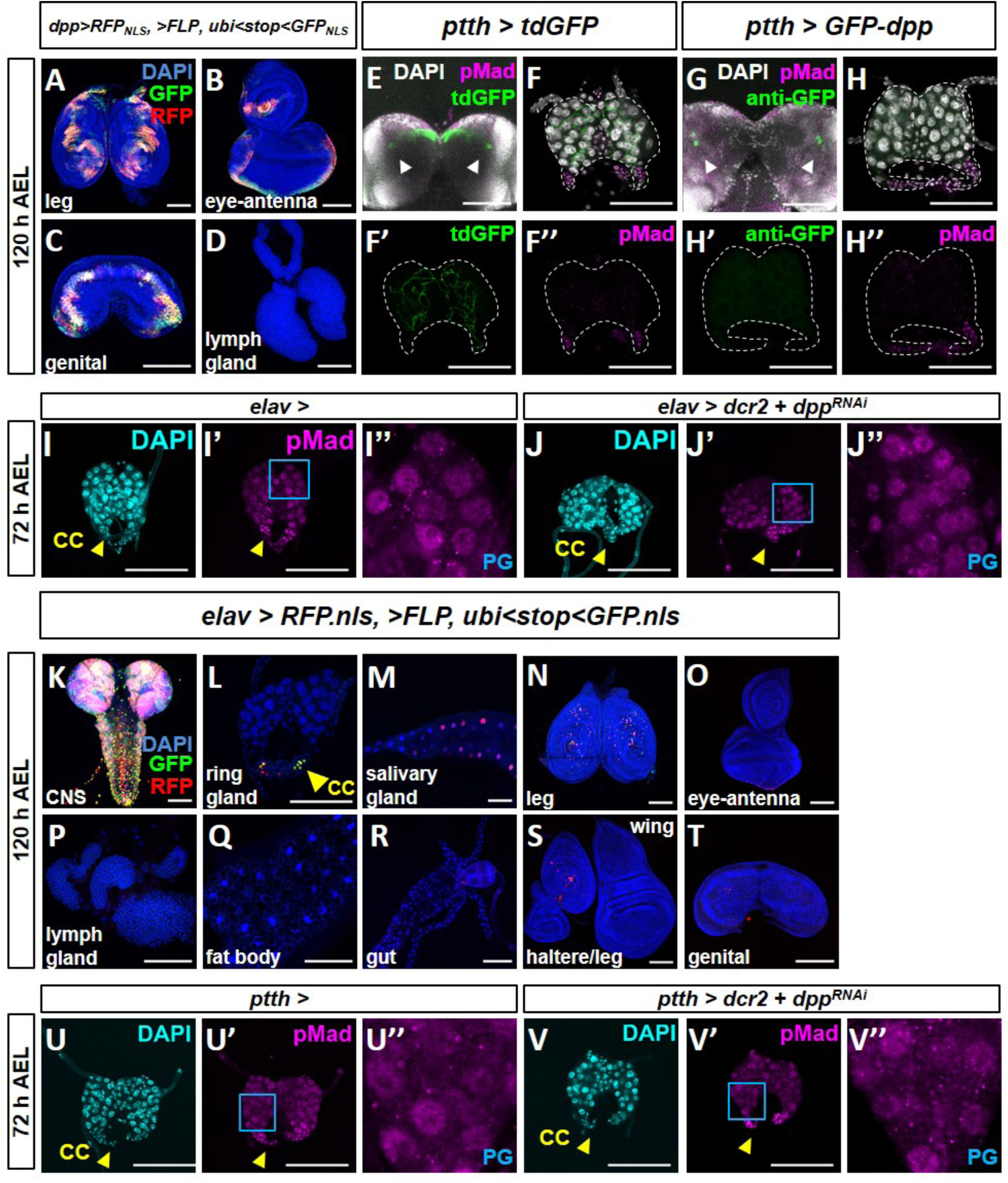
Further characterization of the source of Dpp that reaches the PG in L3 larvae. (**A-D**) Additional images from G-TRACE experiments (*16*) to characterize expression of *dpp-Gal4*. Expression is observed in leg discs (**A**), eye-antennal disc (**B**), genital disc (**C**) but not in the lymph gland (**D**). A previous study (*37*) reported the expression of a *dpp-lacZ* reporter in the CA of the ring gland. In that same study, *dpp* expression in the ring gland, measured using quantitative RT-PCR from total RNA prepared from ring glands, was reported to increase in late L3, peaking in wandering L3 larvae. These observations are not easy to reconcile with our finding that Dpp signaling in the PG decreases during L3. We have not observed any pMad gradient in the PG based on the distance of cells from the CA. We have also not observed any current or past expression of *dpp-Gal4* in the ring gland using G-TRACE (Fig. 3T) or *UAS-tdTom* (Fig. 3U). *LG-dpp, lexOpmCDB::GFP* is also not expressed in the ring gland (Fig. S2F). Since *dpp-Gal4* is not expressed in the ring gland and *dpp-Gal4 UAS-dcr2, UAS-dpp*^*RNAi*^ can reduce pMad expression in the PG, the source of Dpp that acts on the PG is unlikely to be in the ring gland itself. (**E, F-F”, G, H-H”**) Effect of expressing *GFP-dpp* in the PTTH neurons. Expression of *tdGFP* using *ptth-Gal4* which labels cell membranes (**E, F, F’**) shows axons that innervate the PG but does not result in nuclear pMad in the PG. GFP is visualized using GFP fluorescence. Expression using *GFP-dpp* using *ptth-Gal4* (**G, H, H’**) does not result in delivery of GFP-Dpp to the PG (**G, H, H’**) or an increase in nuclear pMad in the PG (**H”**). In (**G, H,-H”),** anti-GFP antibody was added to detect even low levels of GFP expression. Arrowheads indicate PTTH neurons. (**I-I’’, J-J’’**) Effect of *elav-Gal4, UAS-dpp*^*RNAi*^ in ring glands dissected at 72 h AEL. Nuclear pMad is detected both in *elav-Gal4* (**I-I’’**) and *elav-Gal4, UAS-dcr2, UAS-dpp*^*RNAi*^ (**J-J’’**) cells in the PG. (**K-T**) Characterization of expression of the *elav-Gal4* driver using G-TRACE. Note expression in the brain and ventral nerve cord (**K**), CC cells in the ring gland (**L**), salivary gland (**M**), leg disc (**N**) and eye-antennal disc (**O**). No expression in lymph gland (**P**), fat body (**Q**), gut (**R**), wing and haltere discs (**S**), and genital disc (**T**). (**U-V”**) Effect of *ptth-Gal4, UAS-dpp*^*RNAi*^ in ring glands dissected at 72 h AEL. Nuclear pMad is detected both in *ptth-Gal4* (**U, U’, U’’**) and *ptth-Gal4, UAS-dcr2, UAS-dpp*^*RNAi*^ (**V, V’, V”**) cells in the PG. Scale bars are 100 μm.

**Supplementary Figure S7:**
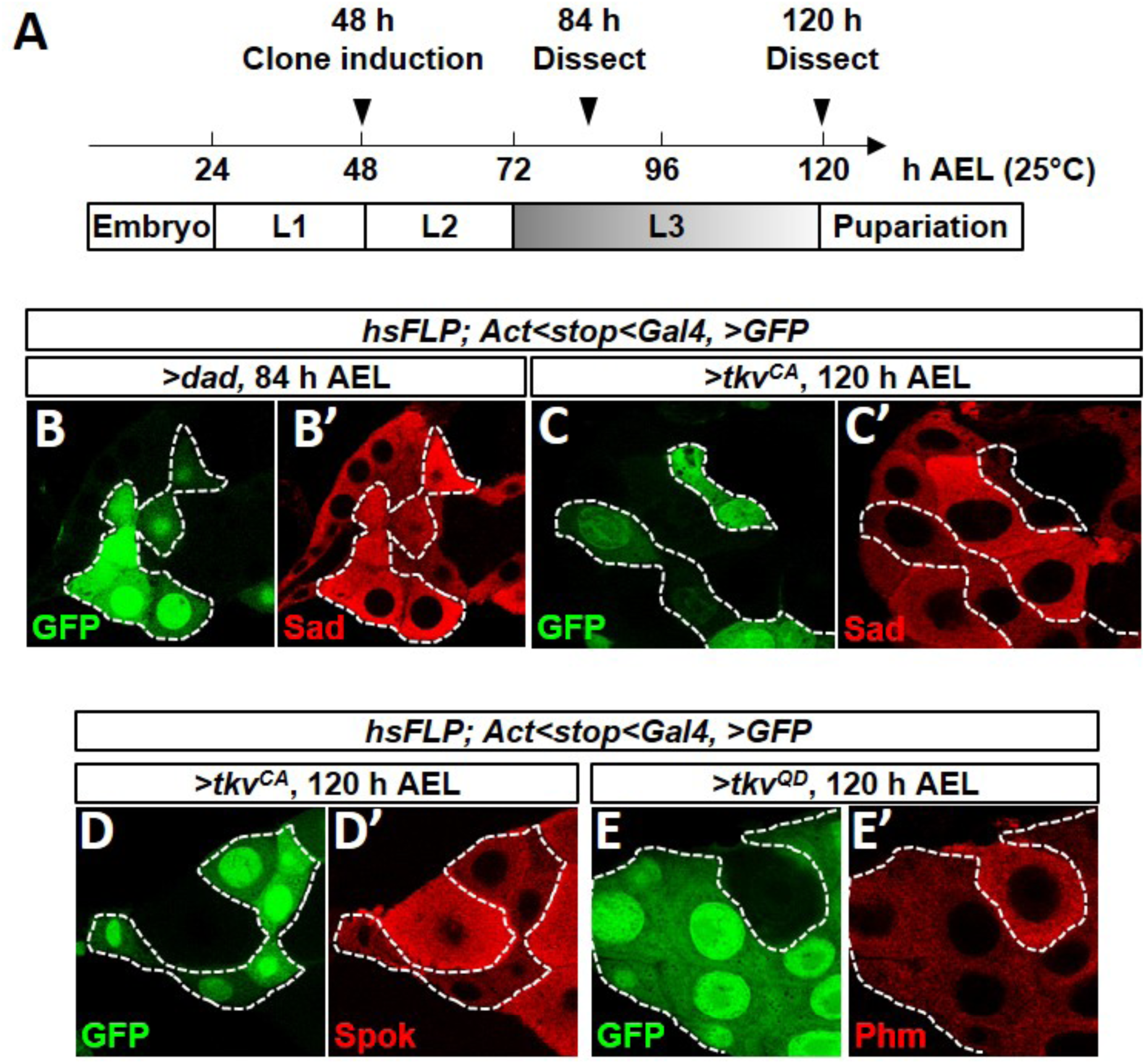
Further characterization of regulation of ecdysone biosynthesis enzymes by Dpp signaling. (**A**) Schematic of heat-shock clone induction experiments. Cultures were maintained at 25°C. FLP-out clones (*act<stop<Gal4*) were induced 48 h AEL. FLP-out clones in the PG show uneven expression of GFP possibly because of asynchrony of or variability in the number of endocycles between the individual polyploid cells of the PG. (**B, B’**) Cell-autonomous reduction in Dpp signaling in 84 h AEL PG cells using *UAS-dad* induces upregulation of Shadow (Sad), although the effect is less obvious than with Disembodied (Dib) (shown in Figure 4). GFP-positive cells express *UAS-dad*. (**C-E, C’-E’**) Cell-autonomous activation of the Dpp pathway in 120 h AEL PG cells by expressing *UAS* transgenes that express an activated form of Tkv (*UAS-tkv*^*QD*^ in E; *UAS-tkv*^*CA*^ in C and D) causes a reduction in levels of Shadow (Sad) (**C, C’**), Spookier (Spok) (**D, D’**), and Phantom (Phm) (**E, E’**). GFP-positive cells express *UAS-tkv*^*QD*^. Once again, the effect on Sad expression is less than with the other enzymes.

**Supplementary Figure S8:**
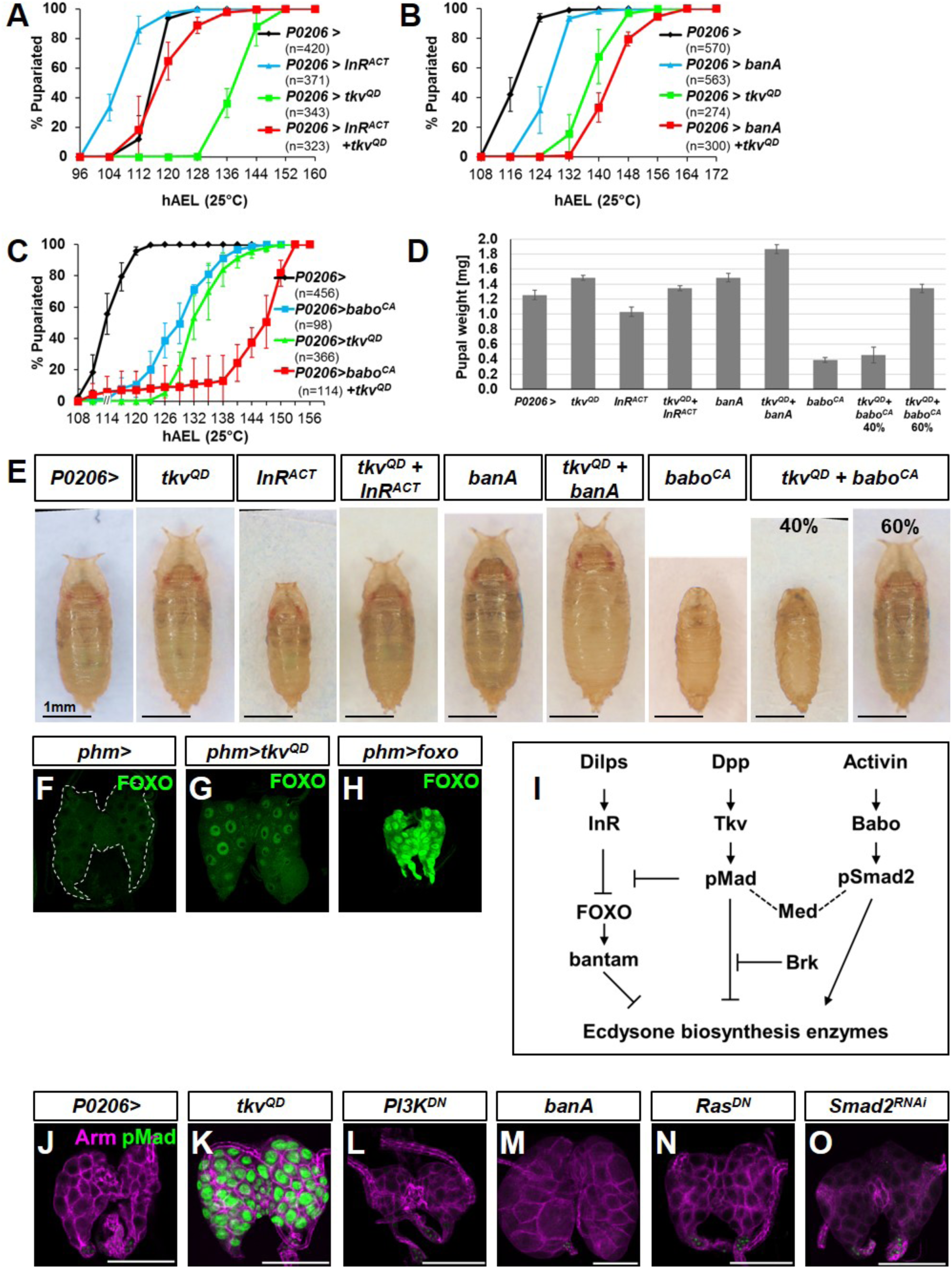
Dpp signaling acts along with other pathways that regulate ecdysteroidogenesis in the PG. (**A-E**) Activation of different signaling pathways in the ring gland using *P0206-Gal4.* (**A, D, E**) Interaction of Dpp signaling with the insulin pathway in the PG. (**A**) *UAS-tkv*^*QD*^ delays pupariation. Expression of an activated form of the insulin receptor (*UAS-InR*^*ACT*^) accelerates pupariation. With concurrent expression of both transgenes, pupariation timing is similar to that of the *P0206-Gal4* driver alone. Pupariation times were: *P0206>* 115.8 ± 0.3 h; *P0206>tkv*^*QD*^ 140.7 ± 7.2 h; *P0206>InR*^*ACT*^ 106.3 ± 1.3 h; *P0206>tkv*^*QD*^ + *InR*^*ACT*^ 117.0.7 ± 3.0 h. Similarly, *P0206> InR*^*ACT*^ pupae are small, *P0206>tkv*^*QD*^ pupae are large and *P0206> InR*^*ACT*^ *+ tkv*^*QD*^ pupae are similar in size or weight to *P0206>* pupae (**D, E**). (**B, D, E**) Interaction of Dpp signaling with *ban*. *UAS-banA* also delays pupariation. Pupariation times were *P0206>* 117.0 ± 1.0 h; *P0206>tkv*^*QD*^ 138.0 ± 2.6 h; *P0206>banA* 125.7 ± 1.5 h; *P0206>tkv*^*QD*^ *+ banA* 143.0 ± 1.7 h. Thus, co-expression of *UAS-tkv*^*QD*^ and *UAS-banA* delays pupariation more than either transgene alone. Since the delay elicited by *UAS-banA* overexpression is less than that obtained with *UAS-tkv*^*QD*^, it is unlikely that *tkv*^*QD*^ delays pupariation exclusively by increasing *ban* expression. Similarly, the increase in pupal size or weight obtained with co-expression of *UAS-tkv*^*QD*^ and *UAS-ban* is greater than the effect obtained with either transgene alone (**D, E**). Note that even though the pupariation delay obtained with *UAS-banA* is less than that obtained with *UAS-tkv*^*QD*^, the effects on pupal size are similar suggesting that *UAS-banA* is more effective than *UAS-tkv*^*QD*^ in increasing growth rate. (**C, D, E**) Interaction of Dpp signaling with Activin signaling. Pupariation times were *P0206>* 114.5 ± 1.3 h; *P0206>tkv*^*QD*^ 129.2 ± 1.5 h; *P0206>babo*^*CA*^ 133.5 ± 3.1 h; *P0206>tkv*^*QD*^ *+ babo*^*CA*^ 148.0 ± 3.0 h. Expression of a constitutively-active form of the Activin receptor (*UAS-babo*^*CA*^) delayed development. However, the larvae appeared delayed in L2 generating small pupae as has been described previously (*7*). With co-expression of *tkv*^*QD*^ and *babo*^*CA*^, approximately 40% of the pupae resembled P0206>*babo*^*CA*^ pupae in appearance and stage-precocious pupariation in L3. The remaining 60% resembled pupae that had gone through the L3 stage and formed pharate adults. The delay in pupariation was close to being additive of the delays caused by either transgene alone. It is possible that the combination of Dpp and Activin signaling leads to an either/or phenotype due to a competition of pMad and pSmad2 with Medea (*22*). **(D)** Pupal weights of all of the conditions shown in panels (A-C). Summary of tests of statistical significance of pairwise comparisons of pupal weight measurements. **(E)** Images of pupae of all the conditions shown in panels (A-C). (**F-H**) FOXO protein visualized in ring glands using anti-FOXO antibody. FOXO is cytoplasmic in *phm>* ring glands (**G**) and nuclear in *phm>FOXO* ring glands (**H**). However, in 20% of *phm>tkv*^*QD*^ ring glands we observed nuclear FOXO (**G**). The reason for the incomplete penetrance of this phenomenon is unclear. A similar frequency of nuclear FOXO was observed when using two different transgenes that encode activated *tkv* and also when using the *P0206-Gal4* instead of *phm-Gal4*. Thus Dpp signaling could impact the insulin pathway upstream of FOXO. (**I**) Model for interaction of Dpp pathway with the InR/FOXO/bantam and Activin signaling. (**J-O**) Ring glands dissected at 120hAEL. (**J**) *P0206>*; (**K**) *P0206>tkv*^*QD*^; (**L**) *P0206>PI3K*^*DN*^; (**M**) *P0206>banA*; (**N**) *P0206>Ras*^*DN*^ (**O**) *P0206>Smad2*^*RNAi*^. Other than *P0206>tkv*^*QD*^, none of these other transgenes, that have each been reported to cause a delay in pupariation (**K-N**), causes nuclear pMad accumulation in the PG indicating that these pathways do not activate Dpp signaling in the PG. Scale bars are 100 μm. Table showing all pairwise comparisons for pupal weights in panel (**D**):

| **Tukey's multiple comparisons test**<br>**(pupal weight of each genotype vs. each other genotype)** | **Summary** |
| --- | --- |
| *P0206>* vs. *P0206>tkv<sup>QD</sup>* | *** |
| *P0206>* vs. *P0206>InR<sup>ACT</sup>* | *** |
| *P0206>* vs. *P0206>InR<sup>ACT</sup>+tkv<sup>QD</sup>* | ns |
| *P0206>* vs. *P0206>babo<sup>CA</sup>* | **** |
| *P0206>* vs. *P0206>babo<sup>CA</sup>+tkv<sup>QD</sup> 40%* | **** |
| *P0206>* vs. *P0206>babo<sup>CA</sup>+tkv<sup>QD</sup> 60%* | ns |
| *P0206>* vs. *P0206>banA* | *** |
| *P0206>* vs. *P0206>banA+tkv<sup>QD</sup>* | **** |
| *P0206>tkv<sup>QD</sup>* vs. *P0206>InR<sup>ACT</sup>* | **** |
| *P0206>tkv<sup>QD</sup>* vs. *P0206>InR<sup>ACT</sup>+tkv<sup>QD</sup>* | ns |
| *P0206>tkv<sup>QD</sup>* vs. *P0206>babo<sup>CA</sup>* | **** |
| *P0206>tkv<sup>QD</sup>* vs. *P0206>babo<sup>CA</sup>+tkv<sup>QD</sup> 40%* | **** |
| *P0206>tkv<sup>QD</sup>* vs. *P0206>babo<sup>CA</sup>+tkv<sup>QD</sup> 60%* | ns |
| *P0206>tkv<sup>QD</sup>* vs. *P0206>banA* | ns |
| *P0206>tkv<sup>QD</sup>* vs. *P0206>banA+tkv<sup>QD</sup>* | **** |
| *P0206>InR<sup>ACT</sup>* vs. *P0206>InR<sup>ACT</sup>+tkv<sup>QD</sup>* | **** |
| *P0206>InR<sup>ACT</sup>* vs. *P0206>babo<sup>CA</sup>* | **** |
| *P0206>InR<sup>ACT</sup>* vs. *P0206>babo<sup>CA</sup>+tkv<sup>QD</sup> 40%* | **** |
| *P0206>InR<sup>ACT</sup>* vs. *P0206>babo<sup>CA</sup>+tkv<sup>QD</sup> 60%* | **** |
| *P0206>InR<sup>ACT</sup>* vs. *P0206>banA* | **** |
| *P0206>InR<sup>ACT</sup>* vs. *P0206>banA+tkv<sup>QD</sup>* | **** |
| *P0206>InR<sup>ACT</sup>+tkv<sup>QD</sup>* vs. *P0206>babo<sup>CA</sup>* | **** |
| *P0206>InR<sup>ACT</sup>+tkv<sup>QD</sup>* vs. *P0206>babo<sup>CA</sup>+tkv<sup>QD</sup> 40%* | **** |
| *P0206>InR<sup>ACT</sup>+tkv<sup>QD</sup>* vs. *P0206>babo<sup>CA</sup>+tkv<sup>QD</sup> 60%* | ns |
| *P0206>InR<sup>ACT</sup>+tkv<sup>QD</sup>* vs. *P0206>banA* | ns |
| *P0206>InR<sup>ACT</sup>+tkv<sup>QD</sup>* vs. *P0206>banA+tkv<sup>QD</sup>* | **** |
| *P0206>babo<sup>CA</sup>* vs. *P0206>babo<sup>CA</sup>+tkv<sup>QD</sup> 40%* | ns |
| *P0206>babo<sup>CA</sup>* vs. *P0206>babo<sup>CA</sup>+tkv<sup>QD</sup> 60%* | **** |
| *P0206>babo<sup>CA</sup>* vs. *P0206>banA* | **** |
| *P0206>babo<sup>CA</sup>* vs. *P0206>banA+tkv<sup>QD</sup>* | **** |
| $P0206>babo^{CA}+tkv^{QD}$ 40% vs. $P0206>babo^{CA}+tkv^{QD}$ 60% | **** |
| $P0206>babo^{CA}+tkv^{QD}$ 40% vs. $P0206>banA$ | **** |
| $P0206>babo^{CA}+tkv^{QD}$ 40% vs. $P0206>banA+tkv^{QD}$ | **** |
| $P0206>babo^{CA}+tkv^{QD}$ 60% vs. $P0206>banA$ | ns |
| $P0206>babo^{CA}+tkv^{QD}$ 60% vs. $P0206>banA+tkv^{QD}$ | **** |
| $P0206>banA$ vs. $P0206>banA+tkv^{QD}$ | **** |

| Summary | P value |
| --- | --- |
| ns | $\geq 0.05$ |
| * | 0.01 to 0.05 |
| ** | 0.001 to 0.01 |
| *** | 0.0001 to 0.001 |
| **** | $<0.0001$ |

